# AFP-R: An Open Resource Dedicated to Antifreeze Proteins

**DOI:** 10.64898/2026.08.21.746139

**Authors:** Wei Liu, Yichi Zhang, Dongxue Xiu, Yangchen Liu, Tinglan Wang, Xiaohui Chai, Hao Qu, Yuze Min, Zhuqing Zhang

**Affiliations:** College of Life Sciences, University of Chinese Academy of Sciences, 100049, Beijing, China; School of Life Sciences, Inner Mongolia University of Science and Technology, No. 7, Arding Street, Kundulun District, Baotou City, Inner Mongolia, 014010, China

**Keywords:** antifreeze protein, resource, database, prediction

## Abstract

Antifreeze proteins (AFPs), lower the freezing point via thermal hysteresis activity and/or ice recrystallization inhibition, playing a crucial role in protecting organisms from freezing damage under sub-zero milieu. This property endows them with promising applications in biomedicine and agriculture, ranging from tissue-organ cryopreservation to the development of frost-resistant crops. However, the lack of comprehensive resources dedicated for AFPs hinders further progress in elucidating their functional mechanisms and advancing their applications. Here, we report AFP-R, an online resource comprising AFP-DB and AFP-Predictor. AFP-DB is a comprehensive database with manually curated proteins bearing experimentally validated antifreeze activity derived from published literature, whereas AFP-Predictor is a sequence-based machine-learning model to identify AFPs. AFP-DB stores diverse AFP-related information, including sequences, structures, post-translational modifications, taxonomy and annotations of antifreeze-activity experimental assays. It now holds 186 entries, 607 sub-entries, and 1444 experimental records. AFP-Predictor, an AFP-identification algorithm built on protein language model ESM2 (Evolutionary Scale Modeling 2), is trained on data in AFP-DB and outperforms several existing models. This work offers a valuable resource for systematically dissecting the mechanisms underlying AFP antifreeze activity and will facilitate their broader applications.

## Introduction

In sub-zero environments, antifreeze proteins (AFPs) are essential for the survival of diverse organisms, such as fish[1], insects[2], plants[3], and microorganisms[4], *etc*. AFPs non-colligatively lower the freezing point of bodily fluids via thermal hysteresis (TH) activity and/or ice recrystallization inhibition (IRI), thereby protecting cells from damage[5]. The prevailing model to interpret the molecular mechanism of AFPs’ antifreeze function is the adsorption-inhibition hypothesis[6], according to which AFPs irreversibly attach to specific ice planes through their ice-binding sites (IBS) and restrian ice growth via the Gibbs-Thomson effect. Owing to these antifreeze properties, AFPs hold great value in multiple fields, ranging from biomedical cryopreservation and agricultural freeze protection to food processing and advanced materials engineering[7].

In recent years, computational studies focused on the identification and prediction of antifreeze proteins have increased substantially [8–15], facilitating the establishment of multiple AFP datasets. Among these resources, the positive dataset containing 481 AFP sequences constructed by Kandaswamy et al. remains the most widely used [11], hereafter referred to as AFP-481. Several recent studies have further expanded the scale of positive training samples for model construction [12–15]. For instance, Nishant Kumar et al. developed the AFProPred algorithm [12] based on an expanded positive dataset comprising 8,134 AFP samples (coined as AFP-8134 herein), which exhibits substantially higher coverage compared with the AFP-481 dataset. Nevertheless, existing datasets used for AFP predictive modeling, including AFP-481, AFP-8134, and other related resources[9,13–15],are predominantly retrieved through retrieving sequences directly by keywords in UniProt (Universal Protein Resource)[16], or collected from the Pfam database[17] then expanding them via BLAST (Basic Local Alignment Search Tool)[18], with only a small fraction supported by solid experimental evidence. This may introduce data noise that impedes the development of reliable AFPs prediction algorithms. To date, a high quality, comprehensive, and experimentally verified resource specifically dedicated to antifreeze proteins is still lacking.

In this work, we present AFP-R, an open-access platform that integrates a manually curated antifreeze protein database (AFP-DB) with a machine learning-based AFP prediction model (AFP-Predictor). All data deposited in AFP-DB were manually mined from 256 peer-reviewed articles via rigorous systematic literature screening. The database provides comprehensive annotations, covering species taxonomy, protein sequence, structural information, mutation types, experimental expression information, antifreeze activity assay annotations, and brief summary. We also performed systematic statistical analyses to characterize the distinct features of curated AFPs compared with general protein sequences. Furthermore, leveraging the high-quality experimentally validated data in AFP-DB, we constructed a series of machine learning models for accurate AFP identification. The optimal model, termed AFP-Predictor, adopts sequence embeddings extracted from the protein language model ESM2[19] for feature presentation and utilizes a multilayer perceptron (MLP) classifie r[20]. Notably, AFP-Predictor achieves outstanding predictive performance and outperforms several existing AFP prediction models [21–23].

## Materials and methods

### Curation of the database AFP-DB

Three public databases―the UniProtKB[16], the PubMed[24] and the Web of Science, were utilized to search and screen eligible publications related to AFPs. All literature data were manually mined and independently validated by at least two researchers to ensure data accuracy. The overall data curation workflow is illustrated in Figure 1.

**Figure 1.**
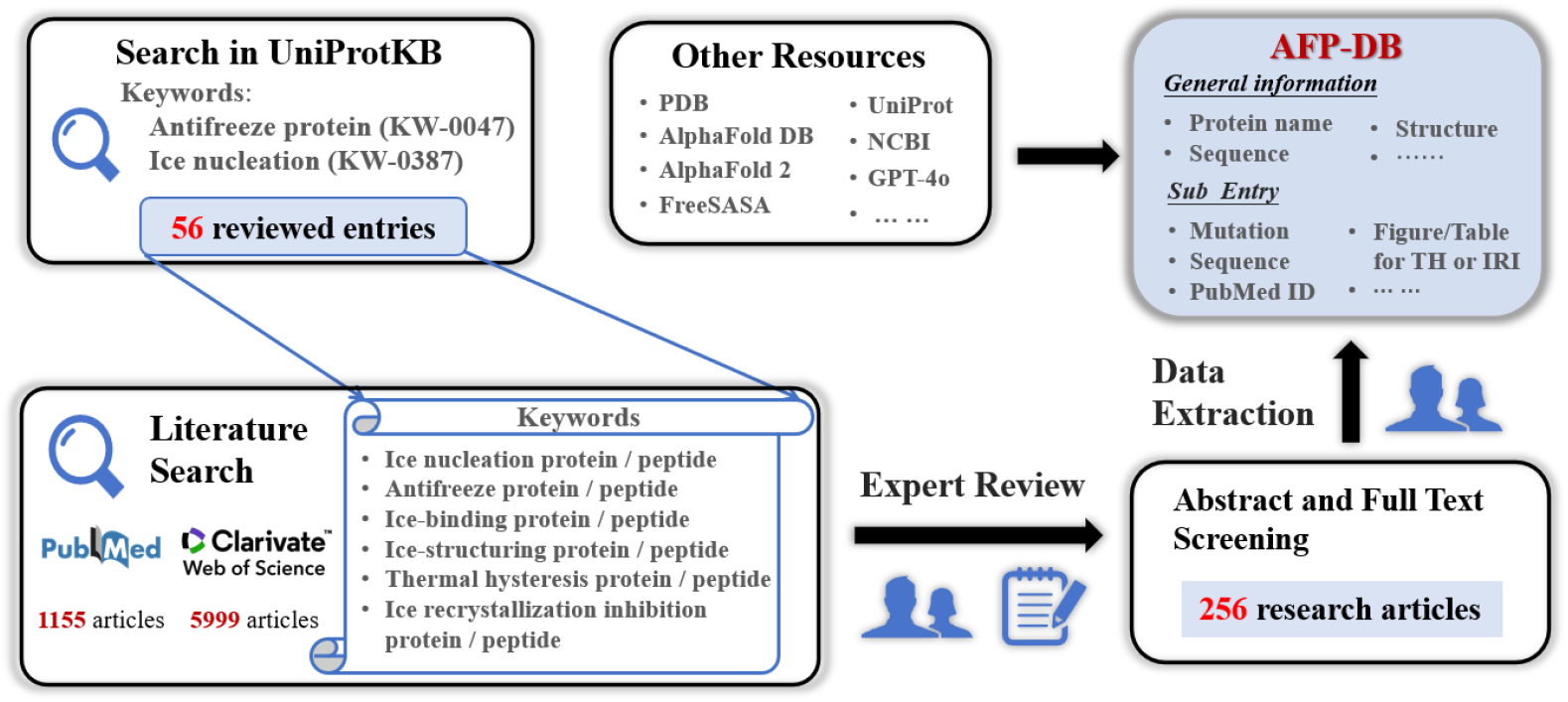
The workflow of AFP-DB construction.

We first searched UniprotKB using keywords “ice nucleation” and “antifreeze protein”, and obtained 56 reviewed entries. After looking through the published literatures corresponding to the 56 reviewed entries, we extended the set of keywords including “antifreeze protein/peptide”, “ice nucleation/peptide”, “ice-binding protein/peptide”, “ice-structuring protein/peptide”, “thermal hysteresis protein/peptide” and “ice recrystallization inhibition protein/peptide”, for further screening publications in the PubMed and the Web of Science databases. The search covered literature up to August 20, 2025, and finally candidate publications including 1,155 ones from PubMed and 5,999 ones from the Web of Science respectively, were collected. These candidate publications were then reviewed by experts. Through abstract screening, followed by full text screening manually, 256 research articles were remained for data extraction, within which review articles, patents and those publications lacking experimental antifreeze activity data were excluded. Papers with antifreeze activity tested only from crude extracts, and those with the antifreeze protein sequence not be able to retrieved, were also excluded. The screening processes and data extraction were conducted independently by at least two experts.

The contents of AFP-DB contain information from the screened 256 articles and that extracted from certain public resources or calculated by related bioinformatics tools. For the former, the following was collected and recorded: (i) General protein information, such as protein name, species and sequence; (ii) More specific information: protein variant and mutation details, modifications, expression tags, *etc*.; (iii) Annotations (legends of the specific figure/table within literatures) of raw experimental data for antifreeze activity assay: thermal hysteresis activity, ice recrystallization inhibition, ice crystal morphology, ice binding plane, *etc*. A brief summary of discussion about protein antifreeze activity in the reviewed publication was recorded with the assistant of artificial intelligence tools GPT-4o and Beijing-YunQue-20230821 under expert supervision. In addition to the information extracted from the literatures mentioned above, we also used other resources such as UniProt and NCBI to complement those data missing in the curated literatures, such as protein sequence, gene name. The three-dimensional (3D) structure of wild-type antifreeze protein in each entry was recorded, which was retrieved from RCSB Protein Data Bank (PDB)[25] or AlphaFold DB[26], or predicted through AlphaFold (version 2.3.2)[27]. As the ice-binding interface of an AFP is crucial for its antifreeze function[28], the solvent-accessible surface area (SASA), and the polar and apolar parts for each naturally occurring or designed AFP were calculated based on 3D structure, through utilizing the prediction tool FreeSASA[29].

The database employs a two-tier hierarchical framework for organizing curated AFPs data, consisting of a primary tier “Entry” and a secondary tier “Sub_Entry”. Each entry represents a unique natural protein, defined by its specific source species and sequence, and generally corresponding to a unique UniProt ID (some entries have no UniProt ID, such as those designed proteins). In each entry, general information including species, protein name, gene name, wild-type sequence, structure *etc*., was recorded. Each entry contains many subentries, with each of them corresponds to the wild type or a specific variant or a designed AFP. Specific information (such as PubMed ID, sequence, annotations of antifreeze activity characterization, *etc*.) was annotated in subentries.

### Datasets for antifreeze proteins prediction

Taking advantage of the curated data from AFP-DB, we built positive datasets for models to identify antifreeze proteins. Positive samples from AFP-DB were filtered by the following criteria: (i) Excluding AFPs with conflicting experimental results across different studies; (ii) Excluding AFPs with fewer than 10 amino acids or containing non-standard amino acids; (iii) Removing redundant AFPs by applying CD-HIT[30] with a threshold of 90% similarity. Finally, 153 AFP sequences were remained, which were split into training set (100 sequences), validation set (25 sequences) and internal test set (28 sequences).

Negative samples were retrieved from the UniProtKB/Swiss-Prot database and filtered by the following criteria: (i) Protein sequences annotated with antifreeze-related keywords (“antifreeze protein”, “ice-binding protein”, “thermal hysteresis”, *etc*.) were excluded; (ii) Protein sequences with amino acid number out of the range 10–2,000 or containing non-standard amino acids were excluded; (iii) Removing redundant proteins by applying CD-HIT[30] with a threshold of 90% similarity. Finally, 333618 non-AFP sequences were obtained. To alleviate the imbalance between positive and negative samples, 653 sequences were randomly selected to build negative dataset from the filtered non-AFP sequences. Among the negative samples, 28 sequences were used for internal test (combined with 28 AFPs in the internal positive test set to form a complete test set with a 1:1 ratio). The remaining 625 sequences were split evenly into 5 folds. The 125 sequences within each fold were divided into a negative training subset and a negative validation subset at a 4:1 ratio.

Due to the limited number of AFPs in AFP-DB, we built an external test set with positive samples retrieved from UniProtKB by antifreeze-related keywords (including “IRI”, “AFP”, “antifreeze polypeptide”, “Antifreeze protein”, “Ice-structuring protein”, “Thermal hysteresis”, “Anti-freeze protein”, “Ice recrystallization inhibition protein”, “Ice-binding protein” and “Ice nucleation”, with “OR” logic), at same time annotated with “experimental evidence at protein or transcript level”. These samples then were filtered by excluding overlapped sequences with the positive training, validation and internal test data, as well as with positive samples in AFP-481[11] and AFP-8134[12] (through CD-HIT with a threshold of 90% similarity). It should be noted that these positive samples were not screened as strictly as those in training sets, meaning that they are “potential AFPs”. Through this strategy, 164 positive samples were retained. An equal number of negative samples were randomly selected from 333618 non-AFP sequences as mentioned above, with those 653 sequences in negative training, validation and internal test sets excluded.

### Model construction and evaluation for identifying antifreeze proteins

Based on the data in AFP-DB, we aimed to construct models to identify AFPs from general proteins. Taking a protein sequence as input, we adopted two strategies to extract protein features. One is to characterize protein with explicit features including Amino Acid Composition (20 dimensions), Amino Acid Index (using Z-scales, 25 dimensions)[31], Dipeptide Composition (400 dimensions), Blocks Amino Acid Substitution Matrices (400 dimensions)[32], Normalized Moreau-Broto Autocorrelation and Complexity Region(154 dimensions)[33], as well as Protein Secondary Structure (3 dimensions), which were designed mainly based on the experiences of existing AFP classifiers[12, 34]. The other is utilizing the Evolutionary Scale Modeling 2 (ESM2) protein language model[19] with 650M parameters, and embeddings from the final (33rd) transformer layer (1,280 dimensions) were extracted, to present protein with implicit features. Followed the presentation of protein features (a 1,002-dimensional vector for explicit features and a 1,280-dimensional vector for implicit features), four classical machine learning classifiers were adopted independently–logistic regression (LR)[35], support vector machine (SVM)[36], random forest (RF)[37], and multilayer perceptron (MLP)[20]. Then each of the eight models gives a predicted score for a specific input protein sequence to identify whether the corresponding protein is an AFP. The whole framework of the AFP identifying models in this study is shown in Figure 2.

**Figure 2.**
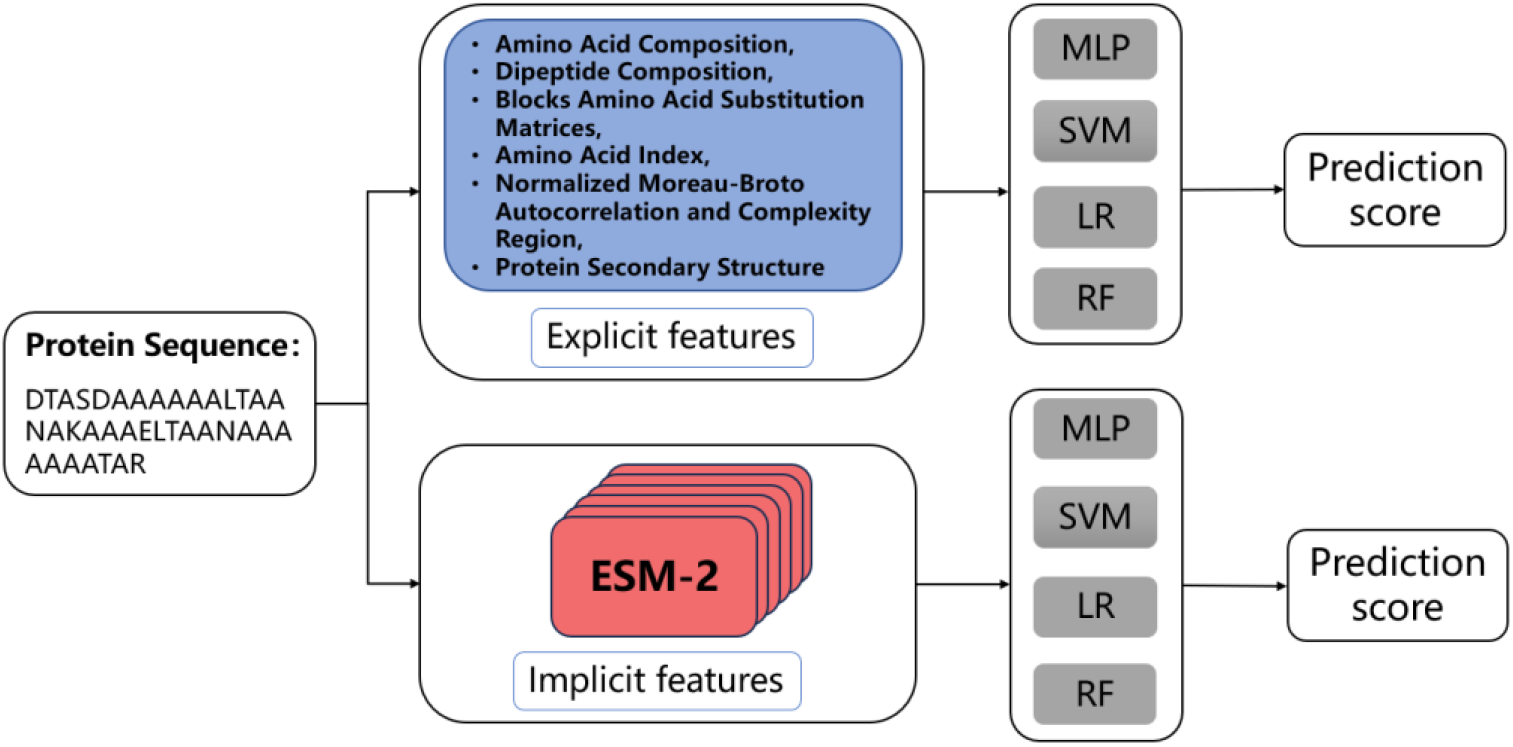
The framework of AFP identifying models constructed in this study.

Due to the number of positive samples is much smaller than that of negative samples, the training strategy analogous to a five-fold cross-validation was employed. Each of the 5-fold negative training/validation dataset was combined with the whole positive dataset to form a complete training/validation set, therefore resulting in five independent sub-models. During the training process, hyperparameters were optimized using the Optuna framework[38] with 100 trials per model, the area under the receiver operating characteristic curve (AUROC) on the validation set was adopted as the optimization objective.

Model evaluation parameters include: AUROC, accuracy (ACC), specificity (SP), sensitivity (SN), F1 score, and Matthews correlation coefficient (MCC)[39], with the corresponding formulas (1)-(5) shown in the following. The classification threshold was determined by maximizing the Youden index (J)[40] (formula (6)). Within these formulas, TP, TN, FP, and FN denote the numbers of true positive, true negative, false positive, and false negative samples, respectively. To evaluate the performance of each type of the studied models, outputs from the five sub-models were averaged.

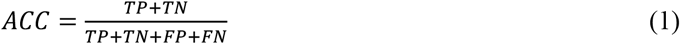

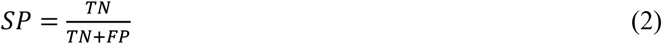

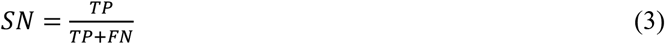

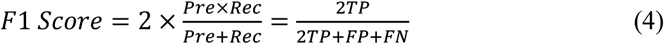

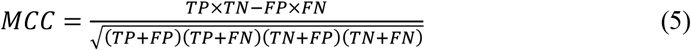

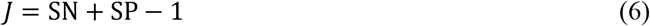

### Implementation of web site for AFP-R

AFP-R is freely accessible online. AFP-DB was built based on Django framework (version 5.1.3), which adopts MTV (Model–Template–View) pattern. The front end of the website was implemented using HTML, CSS and JavaScript, and the back end using Python. MySQL Community Server (version 5.7.19) was used to achieve data storage and management. The implementation was completed under PyCharm environment. The online server AFP-Predictor was mainly developed in Python.

## Results and Discussion

### AFP-DB overview and classification

AFP-DB currently contains 607 subentries which are organized into 186 entries covering curated naturally occurring or engineered AFPs from 256 published articles. All the data were classified into seven categories according to biological source – Fish, Insect, Plant, Bacteria, Diatom, Fungi, and Others, where “Others” means that the AFPs in this category are derived from fusion proteins, enzymatically hydrolyzed short peptides or artificially designed ones. The summary of this classification shown in Table 1 exhibits that except “Others”, the number of entries and that of sub_entries in the “Fish” category are the largest. The other classification according to experimental methodology used to evaluate antifreeze activity is summarized into Table 2. Each experimental record corresponds to an assay documenting the antifreeze activity or inactivity of a given protein, as reported in an individual study. Table 2 shows that the record number of thermal hysteresis activity, which is a critical parameter to quantitatively characterize antifreeze activity of a given AFP, is the largest. These data provide the potential basis for systematic investigation in understanding the relationship between AFP sequence and structure with antifreeze activity.

**Table 1.**
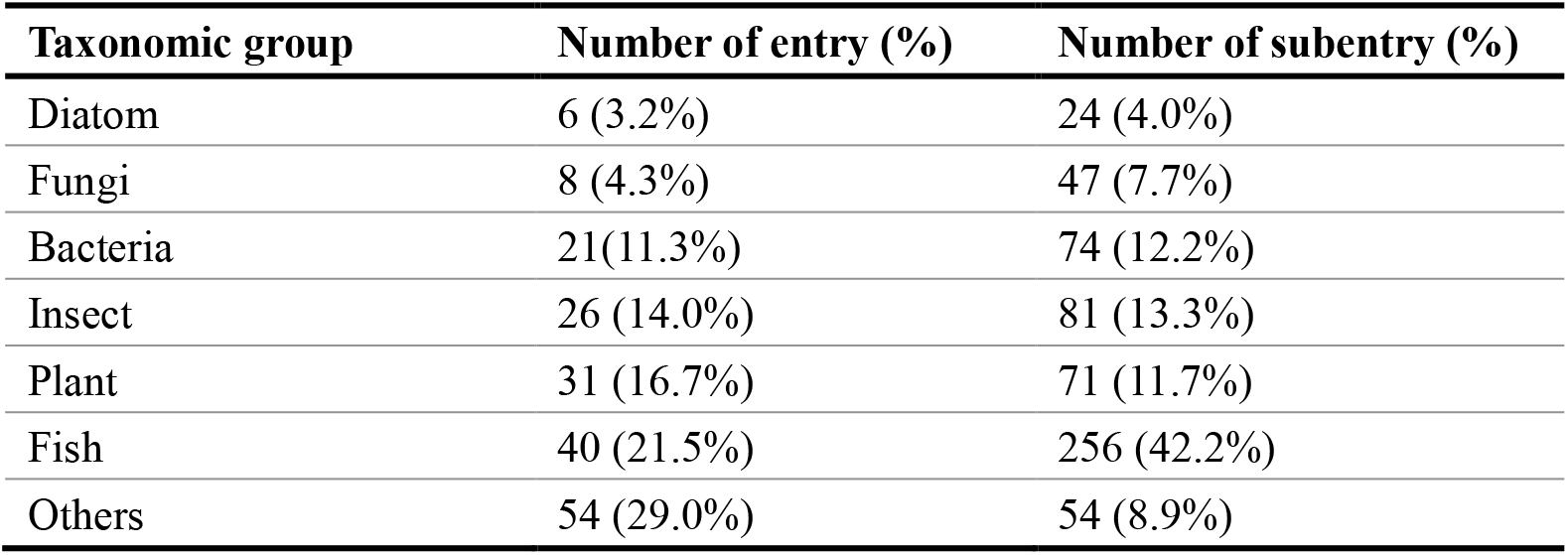
Classification summary of entries according to biological source.

| **Taxonomic group** | **Number of entry (%)** | **Number of subentry (%)** |
| --- | --- | --- |
| Diatom | 6 (3.2%) | 24 (4.0%) |
| Fungi | 8 (4.3%) | 47 (7.7%) |
| Bacteria | 21(11.3%) | 74 (12.2%) |
| Insect | 26 (14.0%) | 81 (13.3%) |
| Plant | 31 (16.7%) | 71 (11.7%) |
| Fish | 40 (21.5%) | 256 (42.2%) |
| Others | 54 (29.0%) | 54 (8.9%) |

**Table 2.**
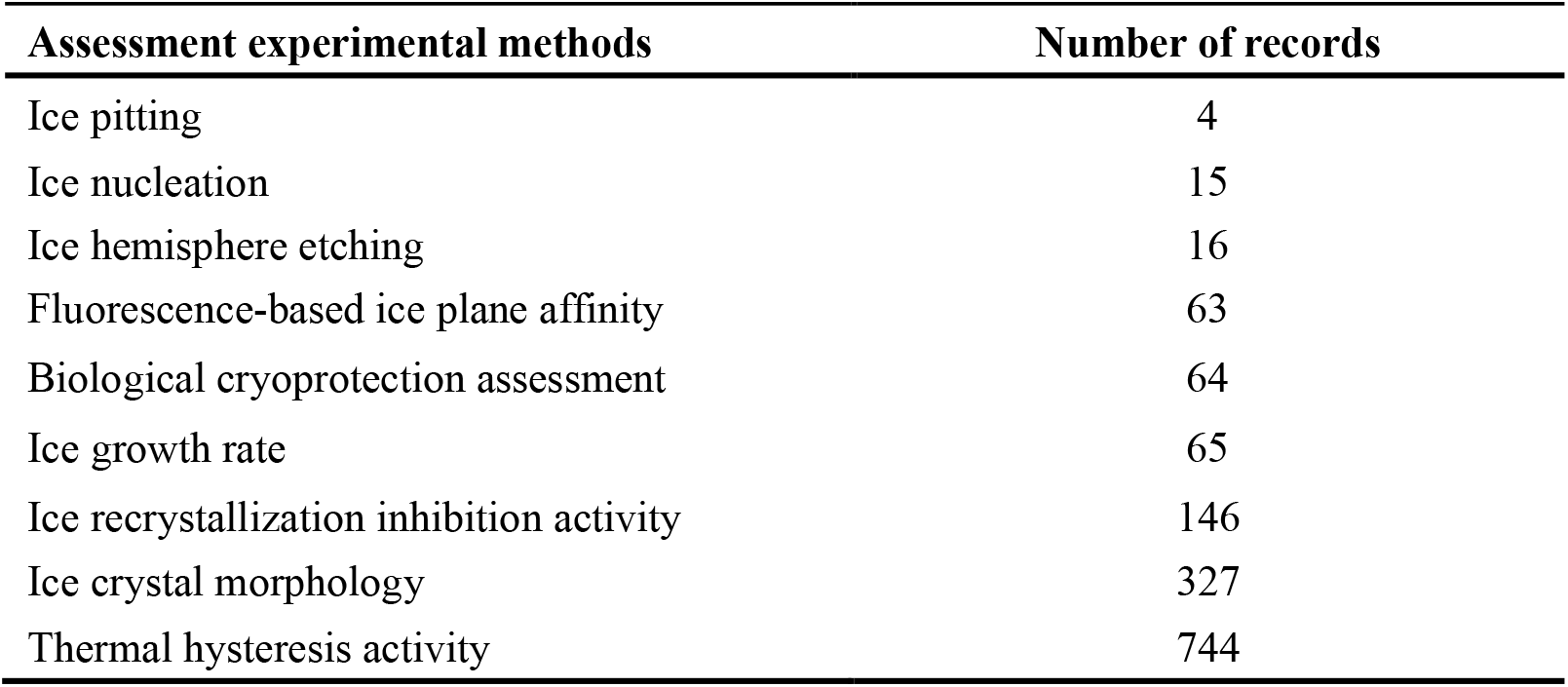
Classification summary of experimental records for antifreeze activity assessment.

### Data statistics of AFP-DB

We used the 132 natural AFPs (excluded 54 entries belonging to the “Others” category) to analyze their characterizations in the following several aspects: (1) The sequence length distribution shown in Figure 3A indicates that most of natural AFPs consist of amino acids with the number less than 600, within which the AFPs from bacteria have the notably longer sequence than others, at the same time with the largest standard deviation (Figure 3B). (2) Amino acids abundances in AFPs across six taxonomic groups are displayed in Figure 3C. Comparing with four representative proteomes (*D. carota, P. americanus, T. molitor* and *Colwellia sp*), alanine, threonine, and glycine exhibit evidently more abundant, while leucine, phenylalanine and arginine show obvious deficiency. This amino acid preference implies the different role of amino acids in antifreeze function of AFPs. (3) Using the bioinformatics tool Foldseek[41] combined with CATH (Class, Architecture, Topology, Homologous superfamily) domain annotations[42], all the AFPs were classified according to their structural features. The results in Figure 3D show that the majority of AFPs falls into classes of “Mainly Beta” and “Alpha Beta”, with lower fraction into “Mainly Alpha” and “Special”. It suggests that the β-sheet structure feature may be generally more apt to offer surfaces to recognize and bind ice crystal plane. (4) Due to the foundation of antifreeze activity is ice binding, we measured both apolar and polar solvent-accessible surface area (SASA) for each AFP through FreeSASA[29] based on its experimental determined otherwise predicted structure, then calculated the ratio between them. The averaged ratio values of AFPs polar/apolar SASA for six taxonomic groups, as well as that of corresponding proteomics (similar species) for each of them are shown in Figure 3E. It is clear that except the taxonomic group “Plant”, the averaged ratios show lower values for AFPs than their corresponding proteomics, suggesting the apolar accessible surface may play crucial function in antifreeze activity.

**Figure 3.**
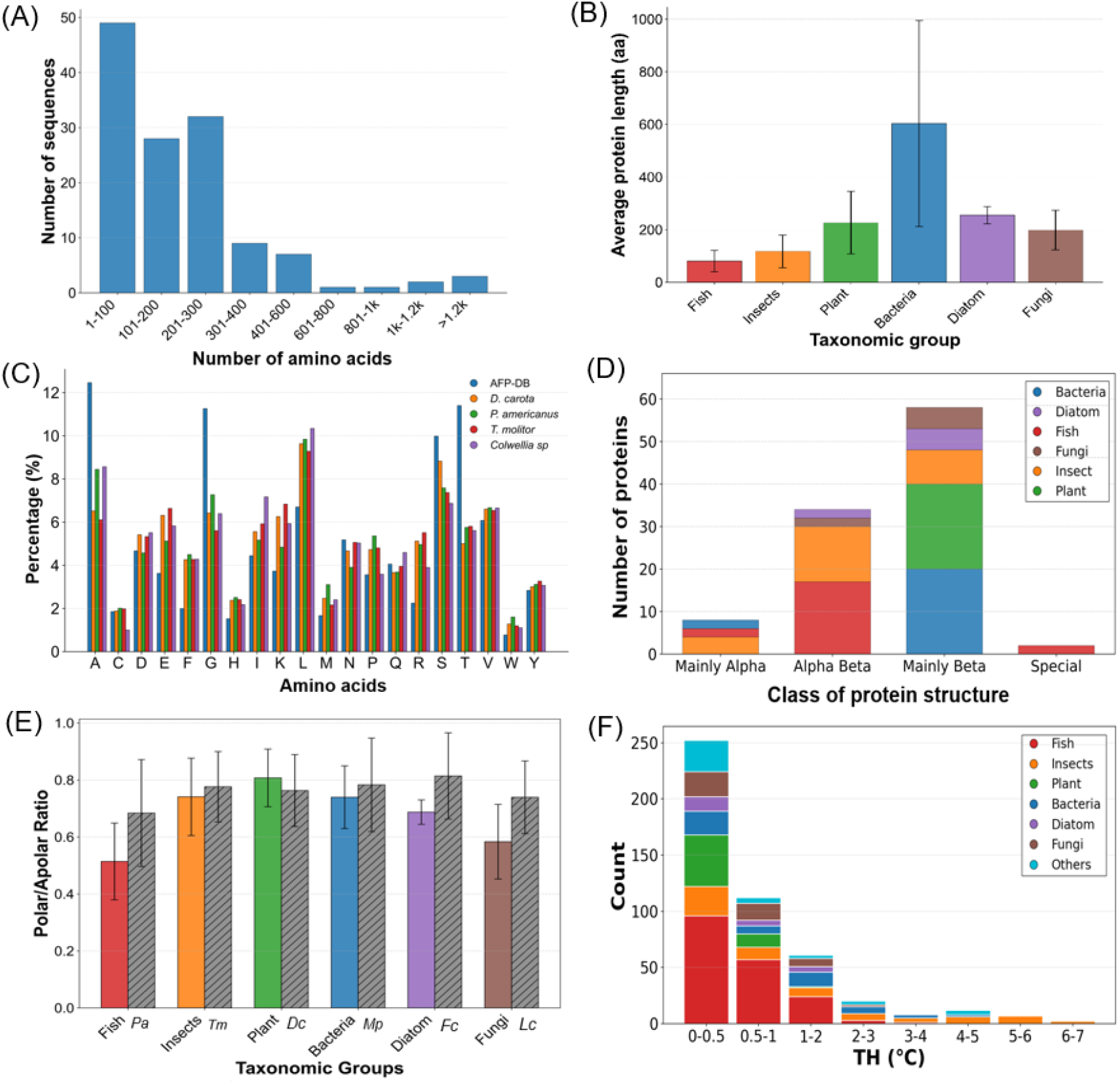
The statistical analysis of the data in AFP-DB. (A) Distribution of sequence length. (B) Averaged sequence length for AFPs from different taxonomic groups. (C) Abundancy of amino acids comparing with proteomics (*D. Carota, P. americanus, T. molitor* and *Colwellia sp*). (D) Distribution of different protein structure classes. (E) Average Polar/Apolar SASA ratio for different taxonomic groups comparing proteomics (*Pa*-Pseudopleuronectes americanus; *Tm*-Tenebrio molitor; *Dc*-Daucus carota; *Mp*-Marinomonas primoryensis; *Fc*-Fragilariopsis cylindrus; *Lc*-Leucosporidium creatinivorum). In (A)-(E), data from “Others” category in AFP-DB were excluded. (F) Distribution of thermal hysteresis (TH) values for all AFPs in AFP-DB.

The thermal hysteresis value recorded in the screened literatures were extracted and the distribution for all AFPs in AFP-DB is shown in Figure 3F. It shows that the thermal hysteresis values measured in experiments are all lower than 8°C, within which the larger values (more than 5°C) are mostly from insect species, while the values from Fish, Plant and Fungi are mostly smaller than 3°C. Further investigation on the thermal hysteresis activity combined with AFP structures as well as the survival milieus of species may provide more insights into understanding the molecular mechanism of biological antifreeze activity.

### Performance of AFP-Predictor

By screening data in AFP-DB and those in UniProt to obtain the training/validation datasets, we utilized two types of protein feature presentation–one is explicit presenting experienced features and the other is implicit taking ESM2 embeddings, combining with four classical classifiers (LR, SVM, RF and MLP), and built eight AFP prediction models based on protein sequence (Figure 2). Table 3 displays the six averaged evaluation parameter values from 5 sub models for each model on internal test set. It is apparent that the models based on implicit feature presentation outperform those based on explicit one, which means the pre-trained protein language model ESM2 can capture the key information encoded in AFPs more accurately. In addition, among the four models based on ESM2, it exhibits that the classifier MLP exhibits better performance than others overall, with the four of six parameters (SN, F1, AUROC and MCC) appearing optimal in Table 3.

**Table 3.** Performance of AFPs prediction models in this study on internal test set.

| Model | ACC | SN | SP | F1 score | AUROC | MCC |
| --- | --- | --- | --- | --- | --- | --- |
| LR-Explicit | 0.9179 ± 0.0411 | 0.8929 ± 0.0565 | 0.9429 ± 0.0741 | 0.9160 ± 0.0408 | 0.9699 ± 0.0191 | 0.8403 ± 0.0815 |
| MLP-Explicit | 0.9500 ± 0.0343 | 0.9286 ± 0.0758 | 0.9714 ± 0.0299 | 0.9478 ± 0.0379 | 0.9908 ± 0.0081 | 0.9037 ± 0.0658 |
| RF-Explicit | 0.9107 ± 0.0357 | 0.8714 ± 0.0896 | 0.9500 ± 0.0407 | 0.9054 ± 0.0411 | 0.9819 ± 0.0084 | 0.8286 ± 0.0689 |
| SVM-Explicit | 0.9179 ± 0.0687 | 0.8643 ± 0.1585 | 0.9714 ± 0.0466 | 0.9068 ± 0.0892 | 0.9867 ± 0.0156 | 0.8503 ± 0.1158 |
| LR-Implicit | **<u>0.9972 ± 0.0032</u>** | 0.9679 ± 0.0233 | **<u>0.9929 ± 0.0160</u>** | 0.9429 ± 0.0479 | 0.9690 ± 0.0221 | 0.9379 ± 0.0443 |
| MLP-Implicit | 0.9821 ± 0.0126 | **<u>0.9857 ± 0.0196</u>** | 0.9786 ± 0.0319 | **<u>0.9823 ± 0.0122</u>** | **<u>0.9995 ± 0.0007</u>** | **<u>0.9651 ± 0.0244</u>** |
| RF-Implicit | 0.9714 ± 0.0160 | 0.9643 ± 0.0357 | 0.9786 ± 0.0196 | 0.9710 ± 0.0166 | 0.9980 ± 0.0028 | 0.9438 ± 0.0318 |
| SVM-Implicit | 0.9750 ± 0.0098 | 0.9714 ± 0.0160 | 0.9786 ± 0.0196 | 0.9749 ± 0.0097 | 0.9962 ± 0.0050 | 0.9504 ± 0.0199 |

Note: Underlined bold font numbers mean the largest values for the corresponding parameters.

To further assess the generalizability of these models, we evaluated them on an independent external test set (See “Datasets for antifreeze proteins prediction” section for details). Three existing AFPs prediction methods which are available to run tasks – AFP-LXGB[23], AFP-CKSAAP[21] and AFP-LSE[22] (the threshold score for them set as 0.5), were evaluated on the same test set. The measured six evaluation parameters of these models are shown in Table 4. These results demonstrate that the models based on ESM2 in this study display better performance than those based on explicit feature presentation, also better than existing models of AFP-LXGB, AFP-CKSAAP and AFP-LSE. In addition, among the models based on ESM2, although the classifier SVM outperforms others as the more optimal parameters (ACC, SP, F1 and MCC) obtained from it, the differences between the models on all the parameters (especially for LR-Implicit, MLP-implicit and SVM-Implicit) are actually minute, as shown in the Table 4. It suggests that models based on implicit features are not very sensitive to the choice of machine learning classifier.

**Table 4.** Performance of AFPs prediction models on external test set.

| Model | ACC | SN | SP | F1 score | AUROC | MCC |
| --- | --- | --- | --- | --- | --- | --- |
| LR-Explicit | 0.9482 | 0.9268 | 0.9695 | 0.9470 | 0.9800 | 0.8972 |
| MLP-Explicit | 0.9512 | 0.9390 | 0.9634 | 0.9506 | 0.9840 | 0.9027 |
| RF-Explicit | 0.9146 | 0.8902 | 0.9390 | 0.9125 | 0.9790 | 0.8303 |
| SVM-Explicit | 0.9421 | 0.9207 | 0.9634 | 0.9408 | 0.9884 | 0.8850 |
| LR-Implicit | 0.9756 | **<u>0.9878</u>** | 0.9634 | 0.9759 | 0.9974 | 0.9515 |
| MLP-Implicit | 0.9756 | 0.9756 | **<u>0.9756</u>** | 0.9756 | **<u>0.9991</u>** | 0.9512 |
| RF-Implicit | 0.9573 | 0.9451 | 0.9695 | 0.9568 | 0.9963 | 0.9149 |
| SVM-Implicit | **<u>0.9787</u>** | 0.9817 | **<u>0.9756</u>** | **<u>0.9787</u>** | 0.9988 | **<u>0.9573</u>** |
| AFP-LXGB | 0.8049 | 0.7256 | 0.8841 | 0.7881 | 0.9339 | 0.6176 |
| AFP-CKSAAP | 0.8445 | 0.8231 | 0.8658 | 0.8411 | 0.9206 | 0.6897 |
| AFP-LSE | 0.8994 | 0.9085 | 0.8902 | 0.9003 | 0.9545 | 0.7989 |

Note: Underlined bold font numbers mean the largest values for the corresponding parameters.

Based on the above evaluations across test sets, we named the best-performing model “MLP-Implicit” AFP-Predictor, and further developed a freely accessible online server integrated into AFP-R.

### Web interface of AFP-R

AFP-R features a user-friendly web interface that integrates AFP-DB and AFP-Predictor (https://bio-comp.ucas.ac.cn/AFP/). For the former, users can search, browse, download or submit AFP data directly. For the latter, users can run tasks and get the corresponding results online through a designed interactive web interface. A “Help” webpage is offered to assist users to utilize the resource. The two core modules of search and browse for AFP-DB, as well as AFP prediction via AFP-Predictor are briefly introduced as the following.

### Search data in AFP-DB

The search function in AFP-DB can be accessed on the home page. Two searching types – “by Keywords” and “by protein sequence”, are offered. For the former, the keyword types are shown in a drop-down list of the corresponding window. For the latter, BLASTP is used, and related parameters are allowed to be reset when users select “Advanced”. Both types of search result in a simplified table as shown in Figure 4, which is in similar form with that in the default browse page. Users can access the detailed page through the linkage with Entry ID or Sub_Entry ID in the table, also can download the whole searched results (including detailed information) through the download button on the top-right of resulting page.

**Figure 4.**
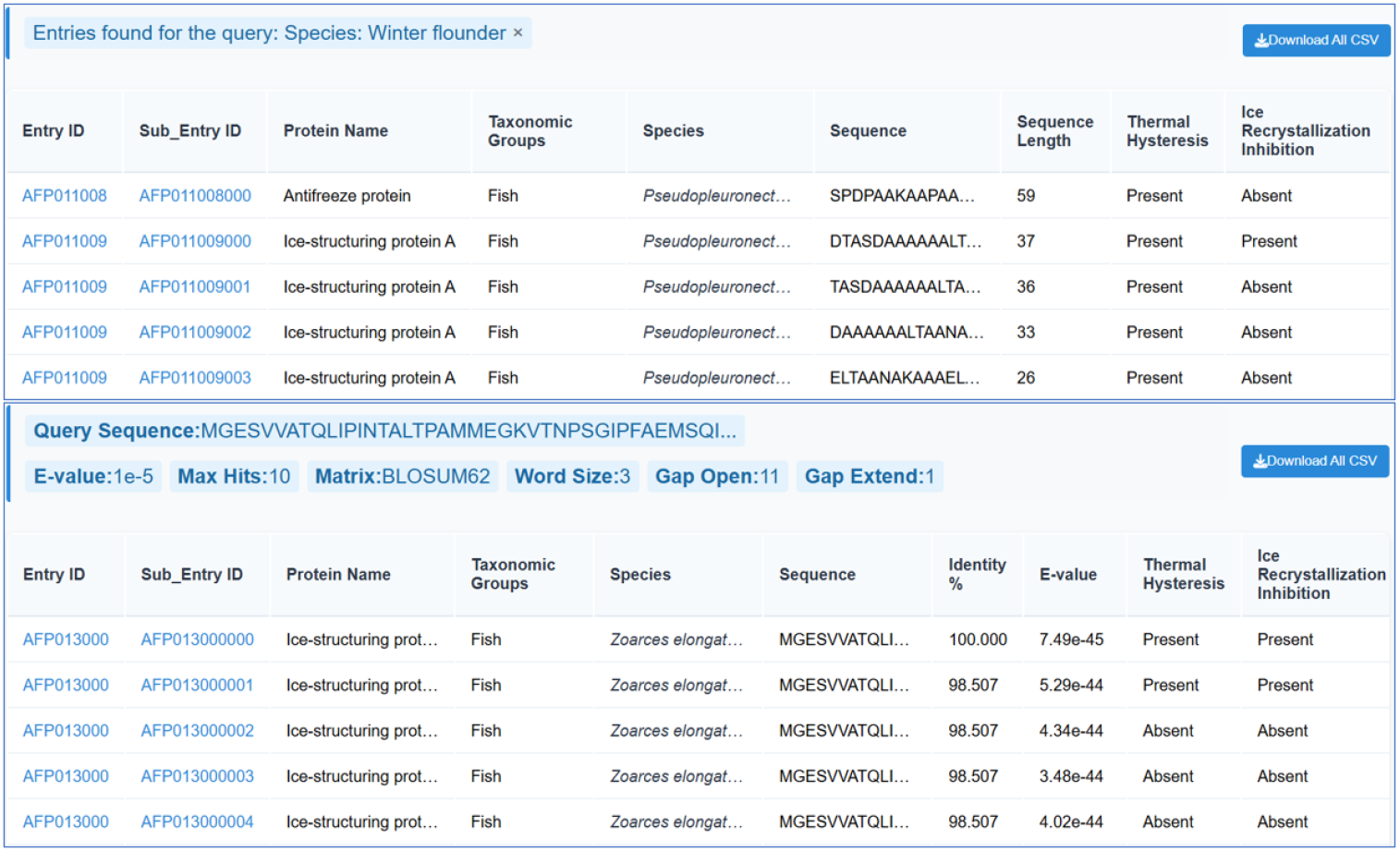
The schematic of searching results “by Keywords” and “by protein sequence”.

### Browse data in AFP-DB

The browse module in AFP-DB includes two layers. The default interface of the first layer contains a categorical navigation bar on the left and a simplified table on the right (in a similar form as that in search resulting page). Two primary classification categories are presented, one based on taxonomic groups (such as Fish, Insect, Plant.) and the other based on antifreeze assay characterization (such as thermal hysteresis, ice recrystallization inhibition activity). The second layer restores the detailed information of entries and subentries (including two tiers as mentioned in “Curation of the database AFP-DB” section). As shown in Figure 5, on each detailed page, the left-hand menu provides navigation to general entry information and subentry information, as well as a download button for the whole page content, and the right-hand side presents the specific contents for the entry and subentries.

**Figure 5.**
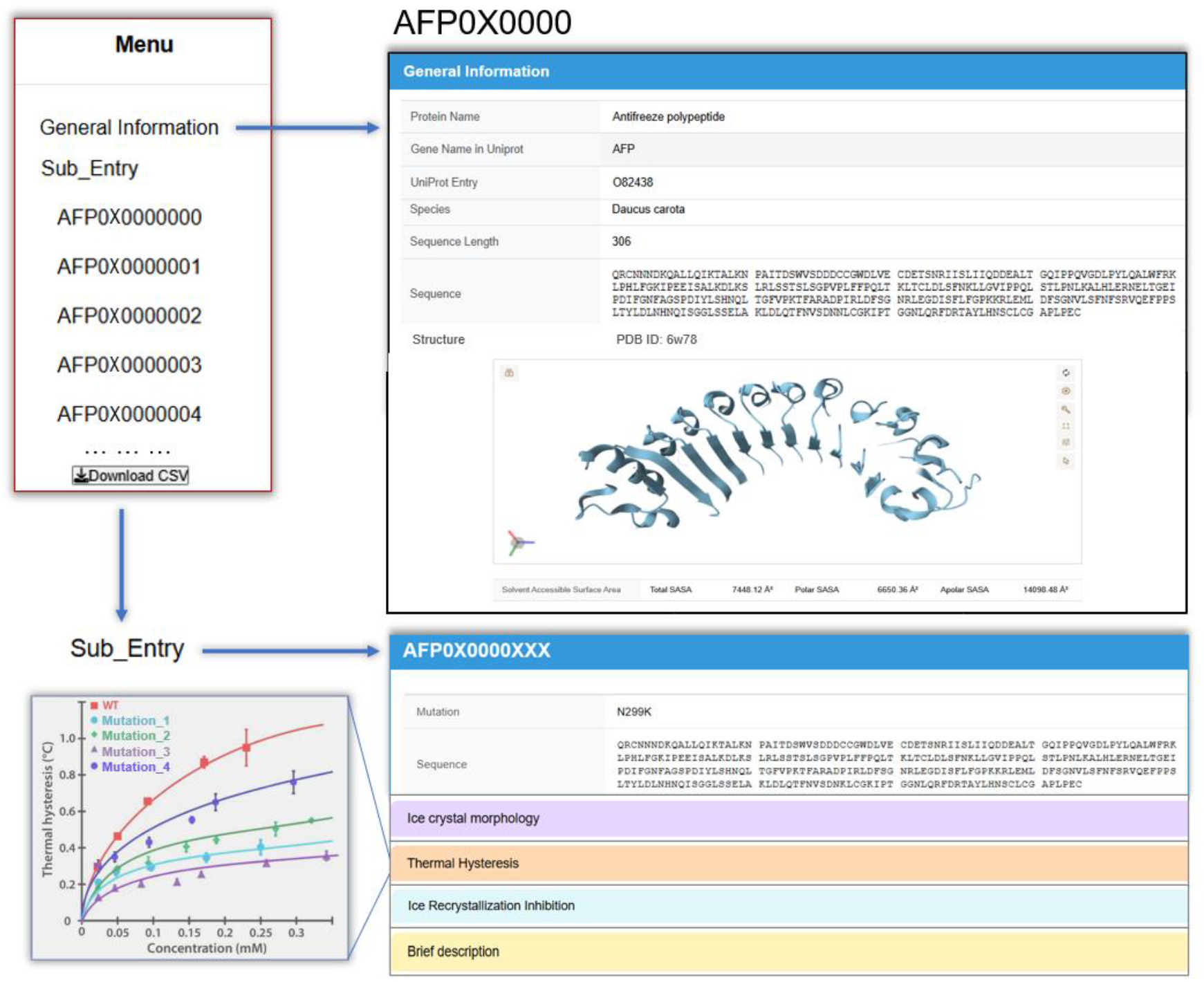
The detailed page for information of entry and subentry.

In AFP-DB, each entry records a unique naturally occurring AFP from a specific biological source or an engineered AFP. The general information of the entry primarily contains protein name, UniProt ID, protein sequence as well as visualization of 3D structure from PDB, AlphaFold DB or AlphaFold2 prediction. The SASA value, as well as its polar and apolar parts based on the structure is also presented in this region. Each subentry records the wild-type or variant sequence, the PMID/DOI, protein name in the literature, mutation type, protein expression information such as expression tag and fusion form, the annotations of the raw antifreeze assay data which include thermal hysteresis activity, ice recrystallization inhibition activity, ice crystal morphology and other related properties, as well as brief summaries of text discussion. Annotations for antifreeze-activity assays document the figure or table legends from the original literature where the experimental data appear, instead of storing the raw figures/tables (such as the lower-left shadowed panel in Figure 5). Users need to access the original publications to view these raw graphics/tables. In addition, for some model AFPs, experimental data might come from different literatures, in this case, more than one annotation record of the same type of antifreeze assay appear in one subentry, with each corresponding PMID/DOI and other related information presented in the specific recording region. At the same time, some raw figures/tables contain measured data for more than one AFP (such as wild type and variants), their corresponding annotations then repeatedly emerge in multiple subentries.

### AFP prediction via AFP-Predictor online

An easy-to-use online web server of AFP-Predictor is provided. The server only requires a protein sequence in FASTA format as input. The outputs include a prediction score and a classification result determining whether the query is a potential AFP, based on a threshold of 0.7796.

## Conclusions

In this work, we constructed AFP-R – an openly accessible resource dedicated to antifreeze proteins (https://bio-comp.ucas.ac.cn/AFP/). It comprises a comprehensively curated database (AFP-DB) and a sequence-based AFP prediction model (AFP-Predictor). The database currently holds 186 entries, 607 subentries, and 1,444 experimental records, covering diverse taxonomic groups as well as engineered AFPs. AFP-DB provides meticulous annotations for all included proteins, especially detailed experimental measurements of their antifreeze activity. AFP-Predictor is an AFP identifier tool built on protein language model ESM2. It is trained on the curated datasets from AFP-DB and outperforms several existing prediction models. AFP-R offers a well-organized, high-quality resource for investigating the sequence-structure-activity relationships of antifreeze proteins and further facilitates their practical applications.

## Acknowledgements

We thank Prof. Zhirong Liu at Peking University and Mr. Shaofeng Liao for their helpful discussion.

## Author contributions

Z.Z. designed research; L.W., Z.Y., X.D., L.Y., W.T., C.X., Q.H. and M.Y. performed research; L.W. and Z.Y. analyzed data; L.W. prepared the original manuscript., L.W. and Z.Z. modified and proofed the manuscript. All authors approved the manuscript publication.

## Data availability

AFP-Resource if freely accessible at https://bio-comp.ucas.ac.cn/AFP/. The data sets for AFP-Predictor construction and evaluation are available in the GitHub repository: https://github.com/ucas-biocomp/AFP-Predictor.

